# Understanding variation in maternal health service coverage and maternal health outcomes among districts in Rwanda

**DOI:** 10.1101/516112

**Authors:** Felix Sayinzoga, Tetui Moses, Koos van der Velden, Jeroen van Dillen, Leon Bijlmakers

## Abstract

**Objective:** To identify factors that explain variations between districts in maternal health service coverage and maternal health outcomes.

**Methods:** Individual key informant interviews and focus group discussions using structured topic lists were conducted in May 2015 in four purposively selected districts.

**Results:** The solidarity support for poor people and the interconnectedness between local leaders and heads of health facilities were identified as enablers of health service utilization. Geographical factors, in particular location close to borders with mobile populations and migrants, and large populations with sparsely distributed health infrastructure, exacerbated by hilly topography and muddy roads were identified as barriers. Shortages of skilled health providers at the level of district hospitals were cited as contributing to poor maternal health outcomes.

**Conclusion:** There is a need to take into account disparities between districts when allocating staff and financial resources in order to achieve universal coverage for high-quality maternal health services and better outcomes. Local innovations such as the use of SMS and WhatsApp text messages by health workers and financial protection schemes for poor patients improve solidarity and are worth to be scaled up.

## Introduction

Worldwide in 2015, an estimated 303,000 women died due to complications of pregnancy or childbirth [1]. Most of them died because of severe bleeding after childbirth, infections, hypertensive disorders or unsafe abortions. Low and middle income countries account for 99% of these deaths, with sub-Saharan Africa alone accounting for roughly two-thirds (201,000 deaths in 2015) [1]. Maternal mortality is one of the health outcomes that typically show very wide gaps between rich and poor populations [1].

The 2014/15 Demographic and health survey (DHS) in Rwanda estimated the maternal mortality ratio (MMR) at 210 deaths per 100,000 live births [2], which is significantly less than the ratios reported in the 2010 DHS (476 per 100,000) and the 2000 DHS (1071 per 100,000) [3,4]. Although everything points to significant improvements, deaths related to pregnancy and childbirth are still too common and will need to be reduced further in order to reach the SDG target MMR of 70 per 100,000 by the year 2030.

Globally, health service coverage rates have increased but many countries are still a long way from universal coverage for most essential reproductive, maternal, newborn and child health (RMNCH) interventions. Furthermore, intra-country variations in service coverage rates are reducing in most countries but the pace is slow. The main impediments to the delivery of high-quality services are a combination of health sector specific and non-health sector drivers [5]. Reliable and timely information is required for effective remedial action at local, regional and national health sector management levels, in order to address intra-country inequalities in maternal health care.

The present study was conducted to obtain an in-depth understanding of the factors that explain variations between districts in maternal health outcomes, including certain emergency obstetric care indicators, and identify the prevailing barriers and enablers to improve maternal health.

## Materials and Methods

### Study setting

We performed a mixed-methods study combining individual interviews and focus group discussions that were conducted in May 2015 in four districts: Bugesera, Gicumbi, Nyagatare and Rwamagana. These districts had a combined population of around 1.65 million people [6–9; see Table 1].

**Table 1:** Demographic and geographic information.

|  | **Bugesera** | **Gicumbi** | **Nyagatare** | **Rwamagana** |
| --- | --- | --- | --- | --- |
| **Province** | Eastern | Northern | Eastern | Eastern |
| **Area** | 1,288 km<sup>2</sup> | 829 km<sup>2</sup> | 1,741 km<sup>2</sup> | 682 km<sup>2</sup> |
| **Population in 2013** | 361,914 | 395,606 | 466,944 | 313,461 |
| **Population density** | 280/km<sup>2</sup> | 480/km<sup>2</sup> | 243/km<sup>2</sup> | 460/km<sup>2</sup> |
| **# of health centres** | 16 | 24 | 21 | 15 |
| **# of hospitals** | 1 | 1 | 1 | 1 |
| **Distance to Kigali** | 35 km | 55 km | 161 km | 60 km |

The districts were chosen on the basis of their performance on a set of key maternal health indicators, which included four service coverage, four service process indicators and four health outcomes, according to data reported in 2013 through the national health formation system (Table 2) [10]. Bugesera district was selected because of its relatively good performance on all three types of indicators. Gicumbi district was chosen because of its good performance on health outcome indicators, but poor performance on service coverage indicators, which seemed rather contradictory. For Rwamanaga it was the reverse: high service coverage rates, but poor health outcome indicators. Nyagatare was included in the study because of its relatively poor overall performance in 2013.

**Table 2:** Selected key indicators for the four districts in 2013 and the national average (rank numbers indicated in brackets, <1> indicating the best performing district, and <30> the worst performing district).*

|  | Bugesera district | Gicumbi district | Nyagatare district | Rwamagana district | National average (range) |
| --- | --- | --- | --- | --- | --- |
| *Service coverage indicators* |  |  |  |  |  |
| Pregnant women with 1<sup>st</sup> antenatal care visit in 1<sup>st</sup> trimester of pregnancy | 51%<br><11> | 34%<br><22> | 46%<br><12> | 69%<br><3> | 45%<br>(15-84) |
| Pregnant women with four or more standard antenatal care visits | 42%<br><9> | 27%<br><20> | 22%<br><24> | 62%<br><3> | 35%<br>(9-74) |
| Deliveries at home | 4.5%<br><17> | 3.1%<br><11> | 9.1%<br><28> | 4.4%<br><15> | 4.5%<br>(0.6-10.3) |
| Women who delivered and attended at least one postnatal care visits | 65%<br><11> | 60%<br><18> | 63%<br><13> | 63%<br><14> | 58%<br>(22-86) |
| *Process indicators* |  |  |  |  |  |
| 2-5 Tetanus toxoid vaccinations received | 66%<br><26> | 67%<br><24> | 59%<br><30> | 70%<br>(18) | 74%<br>(59-100) |
| High-risk pregnancies detected during antenatal care visits | 3.9%<br><n/a> | 9.4%<br><n/a> | 6.1%<br><n/a> | 6.9%<br><n/a> | 9.4%<br>(3.9-32.3) |
| High risk pregnancies referred during antenatal care visits | 76%<br><1> | 20%<br><27> | 27%<br><22> | 26%<br><24> | 43%<br>(7-76) |
| Women with (pre-)eclampsia who received magnesium-sulphate | 77%<br><16> | 28%<br><27> | 80%<br><14> | 71%<br><18> | 95%<br>(0-100) |
| Caesarean sections | 14%<br><n/a> | 11%<br><n/a> | 11%<br><n/a> | 16%<br><n/a> | 14%<br>(3-34) |
| *Health outcomes* |  |  |  |  |  |
| Still births fresh, ≥22 weeks or >500 grams | 0.7%<br><7> | 0.9%<br><16> | 1.2%<br><27> | 1.1%<br><23> | 0.9%<br>(0.4-1.5) |
| Still births macerated, ≥22 weeks or >500 grams | 1.0%<br><17> | 0.8%<br><8> | 1.3%<br><28> | 1.2%<br><26> | 1.0%<br>(0.5-1.5) |
| Reported cases of maternal death in health facilities | 7<br><15> | 3<br><6> | 16<br><25> | 5<br><12> | 269<br>(2-32) |
| Reported cases of maternal death in community | 3<br><24> | 2<br><16> | 6<br><30> | 0<br><1> | 48<br>(0-6) |

\* source: National health management information system (references 6-9 and 10)

### Data collection techniques and tools

All interviews and focus group discussions (FGDs) were conducted in English (by TM) and Kinyarwanda (by a Rwandan research assistant). Interviewees and participants in focus group discussions were allowed to express themselves in Kinyarwanda, English or French. In order to provide structure to the discussions, a topic list was developed, which was used in a flexible manner, with ample room to elaborate on issues that were raised during the interviews.

Sixteen FGDs were conducted (four in each district) with community health workers; nurses working at the district hospital (recruited from the maternity, antenatal clinic and neonatology department); the hospital director together with senior nurse managers from health centres (three facilities, one from each of three sectors chosen randomly per district); and the district director of health together with social affairs officers working at sector level (three sectors per district). In Bugesera district, three additional individual key informant interviews were conducted: one with a technical assistant from one of the development partners, the second with the district hospital director and the third with a senior political leader (the vice-mayor in charge of Social Affairs). All interviews were voice-recorded and afterwards transcribed verbatim (in English) by a research assistant.

### Data analysis

A code manual was compiled inductively, based on the interview topic list with key questions and the verbatim reports of each of the FGD. The actual coding, which involved assigning codes to relevant data segments, was carried out by one of the researchers (MT) with the help of MAXA QDA software (version 2.1.2 for Macs). The coded data were then collated into provisional themes. These were then reviewed by two of the other researchers to verify whether the themes were appropriate in relation to the coded extracts and the entire dataset, before arriving at the final set of themes.

## Results

The study results are presented in two parts, of which the first involves common enablers and barriers across the four districts. The second part presents unique factors that explain variations in performance across the four districts.

### Common enablers and barriers

The increased rates of antenatal care attendance and delivery at a health facility observed over the past decade and – at least in part – the relatively low numbers of reported maternal deaths can be attributed partially to the community meetings that are being organized by local leaders. Community health workers (CHWs), whose role is to educate and sensitize communities about health and other social matters, take part in these meetings and actively discuss their experiences in the field. FGD participants argued that concrete actions such as building health centres so as to bring services closer to the population, ensuring emergency referrals through an efficient ambulance system, and promoting enrolment into community health insurance are being implemented across the country so as to increase geographical and financial access to maternal health services. In addition, the government’s efforts to increase staffing levels, train health workers, motivate them, and establish performance contracts that stipulate zero tolerance of negligence and substandard services were cited as key factors that had a positive influence on service quality.

Respondents across the four districts demonstrated a high level of interconnectedness between actors within the health system and good teamwork between health workers and social welfare officers, particularly in maternal health. This was illustrated with examples such as: mixed teams of medical doctors, nursing staff and administrators from the district hospital conducting weekly supervision tours to health centres within their district; and the monthly coordination meetings, in which staff give an account of their respective activities and performance. This had enhanced transparency of operations and accountability for underperformance, and led to more solidarity across teams. It was enhanced by what some of the respondents termed as ‘a supportive political environment’.

In addition, the *‘rétro-information’* received by health centres on patients they had referred to the district hospital was reported to enhance learning and motivate service providers both at the referring health centre and those at the hospital. The use of short message service technology (rapid SMS) through mobile telephones allowed CHWs to easily contact health centre staff (and vice versa), report on key health indicators, respond to emergencies and mobilize life saving medical care. SMS messages were also being used to remind CHWs who among the pregnant women in their respective villages *(secteurs)* was due for her next antenatal consultation.

> “When we send a rapid SMS, we get the response immediately. If you send a message with a request for help when one of your patients has a problem, you get an ambulance very quickly. This very important for us.”
>
> — (FGD with CHW in Nyagatare district)

All the respondents emphasized the importance of having a district governance system in place that is responsive to local needs. Respondents expressed a high level of trust in their local leaders, particularly in maternal health matters. They sometimes created local by-laws to deal with local problems and generally applied zero tolerance to adverse practices, such as long waiting times for patients or absence from work among health workers. Respondents in all four districts described themselves as ‘highly motivated’ to do a good job and achieve targets, not only for maternal health indicators but also for everything else listed in their performance contracts. These contracts comprise personal and collective targets, which are evaluated periodically. For professional health workers, the salary payments are contingent on their performance, which is evaluated twice a year or every quarter.

There also appeared to be a strong internal motivation and personal drive. This was illustrated by many respondents who emphasized they were keen to serve their communities and claimed to strive for excellence. Some said they would not be satisfied unless the antenatal consultations coverage rate attained 100%. Others were proud not to have had a single case of maternal death in their area for a year or more.

One of the community health workers explained her personal motivation:

> “I feel lucky to be a CHW, because it serves my community, Imana (God) will reward us. I’m proud; it’s not for the money that I’m doing this. Sometimes we sacrifice by neglecting farming or our own household chores.”
>
> — (FGD with community health workers in Bugesera district)

Three distinct operational barriers for providing good quality services were brought up, which all had a financial dimension: partial performance of the community health insurance scheme (*mutuelles de santé*), non-functional medical equipment and inadequacy of water supply at hospitals and health centres. Some of the health workers interviewed were of the opinion that the insurance premiums paid by members of the public were barely sufficient to cover the cost of services that health facilities are expected to provide. Patients who do not have health insurance at all put this system further under pressure.

> “There is a limit to which the cost of services can be covered by the revenues from health insurance premiums. For a delivery, for instance, we use up to ten pairs of gloves; beyond that, the health insurance doesn’t cover the cost.”
>
> — (FGD with nurses, Rwamagana district.)

### Unique factors

While some of the enablers and barriers to delivering quality services and achieving favourable maternal health outcomes are common across the four districts, some factors are unique and seem to explain some of the variation in performance among districts.

The relatively high coverage of maternal health services attendance in Bugesera district can be explained by the solidarity support for poor people to utilise health services through a local scheme that FGD participants termed as “ingobyi”. This scheme can take the form of payment of transport costs or exemption to pay the annual premium for health insurance. It is illustrated by the following quote from one of the FGD:

> “The community contributes 300 francs per week;^1^ if it is a poor family of ten people, we may pay the health insurance premium for six of them. Ingobyi is highly valued in the community and supported by our local leaders.”
>
> — (FGD with CHW, Bugesera district.)

The good performance in coverage indicators in Rwamagana district was explained, at least in part, by the interconnectedness between local leaders and the heads of health facilities, which is facilitated by a WhatsApp group. It is being used by nearly 50 people to exchange information, coordinate activities and support each other on various topics. The group has been critical in responding to logistical challenges and providing advice to its members.

The low maternal health service coverage rates in Gicumbi and Nyagatare districts were attributed to their locations, close to Rwanda’s borders with Uganda and Tanzania, respectively. These two districts have large mobile populations and migrants, from Rwanda itself or from nearby countries. According to the respondents, migrants are often not enrolled in health insurance and they are mostly poor people, which limits their access to care. The problem of mobile populations is illustrated by the following quote:

> “They don’t have permanent settlements; they can be away for three or four weeks before coming back again. This makes follow up very difficult. Some of them don’t even have insurance. This has increased the number of home deliveries in our district.”
>
> — (FGD with CHWs in Gicumbi district)

The same two districts also have relatively large populations, compared to the other two districts included in the study and most other districts in Rwanda, along with sparsely distributed health infrastructure. This is exacerbated by the hilly topography (in Gicumbi) and the poor network of muddy roads (Gicumbi and Nyagatare). It delays the response to requests for emergency ambulance transport and eventually makes it difficult to reduce the number of home deliveries. This, it was argued, limits access to maternal health services.

> “Our district is the biggest and we have only one district hospital. We receive pregnant mothers who have walked long distances; many cannot afford the cost of transport by motor bike.”
>
> — (FGD with CHWs, Gicumbi district)

The size and composition of the health workforce is not adequate compared to the population size and the disease profile. The shortage of clinical specialists at district hospitals was a cause for concern, especially in Nyagatare district which is relatively far away from referral hospitals in the capital Kigali. This also happens to be the district with the largest population.

Respondents in Gicumbi and Rwamagana indicated that having a nursing school gives them extra nurses and midwives who come for practice sessions. They seem to be a bit better off in terms of workload of their nursing staff, as they have the advantage of interns who complement their regular staffing levels. Medical equipment was reported to be in short supply in some of the district hospitals and health centres. In Rwamanaga district, several health facilities experienced shortages of running water, which some respondents considered a health hazard.

## Discussion

This multifaceted qualitative study has identified factors that explain variations between districts in maternal health outcomes, as well as enablers and barriers to improve maternal health. The interconnectedness between local leaders and head of health facilities and the solidarity support for poor people (in one of the districts) were identified as enablers of health service utilisation. Having large mobile populations, high numbers of migrants, and geographical remoteness of sparsely distributed health infrastructure, exacerbated by hilly topography and muddy roads, were identified as barriers. Shortages of health providers skilled at the level of district hospitals were cited as contributors to poor outcomes.

Similar to most other low- and middle-income countries, Rwanda has shaped its national health system according to the framework promulgated by WHO, which is composed of six building blocks [11]. While most of these building blocks are relatively easy to conceptualize, it is not always easy to operationalise and make them work at all levels of the health system. An earlier study has shown that district health managers in Rwanda hold rather strong opinions about what works well in the health sector and what actually makes it work: they considered the dense network of community health workers and the health insurance system as factors that have contributed most to Rwanda’s good achievements [12]. They also indicated the importance of good managerial skills and the culture of continuous monitoring of key indicators. The present study confirms this.

It is uncontested that good governance and leadership are key ingredients of a well functioning health system [13,14]. It includes for instance clarity of direction at the national level, strategic planning, effective oversight and accountability for performance. The present study has shown that this also applies to the local level. FGD participants demonstrated strong commitment to achieving national goals and adhering to national strategies, embodying a great deal of purpose and trust in the activities that they and their colleagues undertake to promote health in their respective districts. They also expressed solidarity with people in society who do not have easy access to health services, and emphasized the importance of intersectoral collaboration and good communication with local leaders. This is what one could refer to as ‘responsiveness’. It is this readiness to respond to local needs and to indications that certain targets might not be met without some extra effort, which might explain the overall increase in health service utilization in Rwanda, observed for already more than a decade [2].

Without a functional health information system, which relies on the timely provision and close monitoring of a selected number of key health indicators, the district-level stakeholders interviewed in this study would not have been in a position to display the vigilance that transpired in the interviews and FGD. The provision of mobile phones to community health workers and the use of rapid SMS to promote antenatal consultations has motivated community health workers to do what is expected of them and it has increased their self-esteem [15].

Adequate health care financing, including funding of all the necessary support services, such as supervision, supply chain management and transportation, are widely considered essential for a health system to thrive [16]. The present study has brought out some challenges at the district level to provide adequate coverage to immigrants and internal migrants who in many cases do not have medical insurance. This explains the low coverage of use of maternal health services in Nyagatare and Gicumbi districts, at least in part. One district (Bugesera) has its own local social solidarity mechanism in place in an effort to ensure equitable access, which has contributed to the high maternal health service coverage rates.

Health workforce adequacy is an issue in Rwanda, as the number of staff with clinical expertise does not always seem to match the requirements [17]. In the case of Nyagatare district, some interviewees and FGD participants did mention that the district was much less popular among health workers than other districts due to its geographical location (far from Kigali, with few opportunities for leisure) compared to the other districts included in the study. The fact that two of the other district hospitals have nursing schools attached to them – which is also the case for Nyagatare district hospital – was cited as a factor that helped alleviate the high workload of nurses and midwives.

The promotion of maternal and child health was seen as a joint responsibility of several Government departments and this has translated into very close, almost day-to-day working relations between civil servants, community health workers and village/sector leaders. The critical role of frontline workers as intermediates between the community and the formal health sector has recently also been confirmed in a study that explored how relationships influence performance in Malawi [18]. In another recent study on health district productivity in Cambodia, Ensor et al. considered the impact of health policy changes on health sector efficiency. Higher efficiency was associated with more densely populated areas and with the presence of health equity funds [19]. While the former may not apply to Rwanda – as virtually the entire country is densely populated – the latter may hold as well.

A limitation of the study is that it did not look in an in-depth manner at quality of care issues. Interviewees did acknowledge that poor quality of care may impact on patients’ demand for services, but they did so mostly by referring to circumstances beyond their own control that affected the quality of care in a negative manner. Neither did the study try to assess whether health workers had the right technical background to provide quality maternal health services. It is worth noting though, that Rwanda does have a mechanism in place to conduct maternal death audits, and the country is currently in the process of institutionalizing ‘near-miss’ audits and confidential enquiries into maternal deaths [20]. Some studies recently reported rather high rates of postpartum haemorrhage and infection, as causes of maternal near miss and death [21,22]. This suggests there is a need for improvement of quality of care at the level of district hospitals, where the majority of deliveries occur.

## Conclusions

Disparities between districts in maternal health outcomes are partly attributed to environmental factors, such as topography, population pressure and migration. The two dimensions over which district managers have control are: (1) the functionality of the district health system, in particular local leadership, health information, funding of support services and health workforce issues; and (2) the extent to which supportive intersectoral activities are being undertaken. There is a need to take into account disparities between districts when allocating staff and financial resources in order to achieve universal coverage for high-quality maternal health services and better outcomes. Further research would be useful to explore the potential of scaling up successful local initiatives, such as the use of SMS and WhatsApp group messages and local financial protection schemes.

## DECLARATION

### Ethics approval and consent to participate

The protocol for this study was approved both by the National Health Research Committee (NHRC/2015/PROT/006) and the Rwanda National Ethics Committee (105/RNEC/2015). Informed consent was obtained from all interviewees.

### Authors’ contributions

FS contributed to the study design, data collection, data analysis, interpretation and writing of the manuscript. TM led the data collection and performed the initial analysis. KvdV and JvD contributed to the study design and writing of the manuscript. LB provided critical intellectual input to the study design, participated in the field work and contributed to data analysis and interpretation and to the writing of the manuscript.

## Acknowledgements

The authors thank the administration and staff of Bugesera, Gicumbi, Nyagatare and Rwamagana districts, especially those who participated in the interviews and focus group discussions : community health workers, nurses working at the district hospital (maternity, antenatal clinic, neonatology department), senior nurse managers from health centres, district directors of health, district hospital directors, social affairs officers working at sector level, and the technical assistant and vice-mayor in charge of Social Affairs in Bugesera district. Special thanks go out to Ms Malka Karangwa and Ms Maartje Kletter for their participation in data collection.

